# Katdetectr: An R/Bioconductor package utilizing unsupervised changepoint analysis for robust kataegis detection

**DOI:** 10.1101/2022.07.11.499364

**Authors:** Daan M. Hazelaar, Job van Riet, Youri Hoogstrate, Harmen J. G. van de Werken

## Abstract

**Background:** Kataegis refers to the occurrence of regional genomic hypermutation in cancer and is a phenomenon that has been observed in a wide range of malignancies. A kataegis locus constitutes a genomic region with a high mutation rate, i.e., a higher frequency of closely interspersed somatic variants than the overall mutational background. It has been shown that kataegis is of biological significance and possibly clinically relevant. Therefore, an accurate and robust workflow for kataegis detection is paramount.

**Findings:** Here we present *Katdetectr*, an open-source R/Bioconductor-based package for the robust yet flexible and fast detection of kataegis loci in genomic data. In addition, *Katdetectr* houses functionalities to characterize and visualize kataegis and provides results in a standardized format useful for subsequent analysis. In brief, *Katdetectr* imports industry-standard formats (MAF, VCF, and VRanges), determines the intermutation distance of the genomic variants and performs unsupervised changepoint analysis utilizing the Pruned Exact Linear Time search algorithm followed by kataegis calling according to user-defined parameters.

We used synthetic data and an *a priori* labeled pan-cancer dataset of Whole Genome Sequenced malignancies for the performance evaluation of Katdetectr and five publicly available kataegis detection packages. Our performance evaluation shows that Katdetectr is robust regarding tumor mutational burden (TMB) and shows the fastest mean computation time. Additionally, Katdetectr reveals the highest accuracy (0.99, 0.99) and normalized Matthews Correlation Coefficient (0.98, 0.92) of all evaluated tools for both datasets.

**Conclusions:** *Katdetectr* is a robust workflow for the detection, characterization, and visualization of kataegis and is available on Bioconductor: https://doi.org/doi:10.18129/B9.bioc.katdetectr

## Introduction

Large-scale next-generation sequencing of malignancies has revealed that a myriad of mutational mechanisms and mutational rates are at play, even within the genome of one tumor cell. It has been shown that cancer genomes can contain regions with high mutation rates, which causes mutations to cluster, i.e., the somatic mutations are found in proximity to one another. This phenomenon was termed kataegis, and its respective genomic location was termed a kataegis locus [1-5].

Since the discovery of kataegis, different computational detection tools using genomic variant data have been developed and are publicly available, including; MafTools [6], ClusteredMutations [7], kataegis [8], SeqKat [9] and, SigProfilerClusters [10]. These packages employ distinct statistical methods for kataegis detection and differ in their ease of use and computational feasibility.

Here, we introduce Katdetectr, an R-based Bioconductor package that contains a suite for the detection, characterization, and visualization of kataegis. Additionally, we have evaluated and compared the performance of Katdetectr to five commonly used and publicly available kataegis detection packages.

## Results

The principle of Katdetectr is to assess the variation in the mutation rate of a cancer genome. To achieve this, Katdetectr starts by importing and preprocessing industry-standard variant calling formats (VCF, MAF, VRanges) (**Figure 1A**). Next, the Intermutation Distance (IMD) is determined, which denotes the distance between variants in base-pairs (**Figure 1B**, see Methods). Unsupervised changepoint analysis is performed, using the IMD as input, which results in detected changepoints. The changepoints, which denote the points at which the distribution of the IMD changes, are used to segment the genomic sequence. Finally, segments are annotated and labeled as a putative kataegis locus if a segment fits the user-defined settings: the mean IMD of the segment ≤ *IMDcutoff* and the number of variants in the segment ≥ *minSizeKataegis*. The IMD, segmentation, and detected kataegis loci can be visualized by Katdetectr in a rainfall plot (**Figure 1C**).

**Figure 1:**
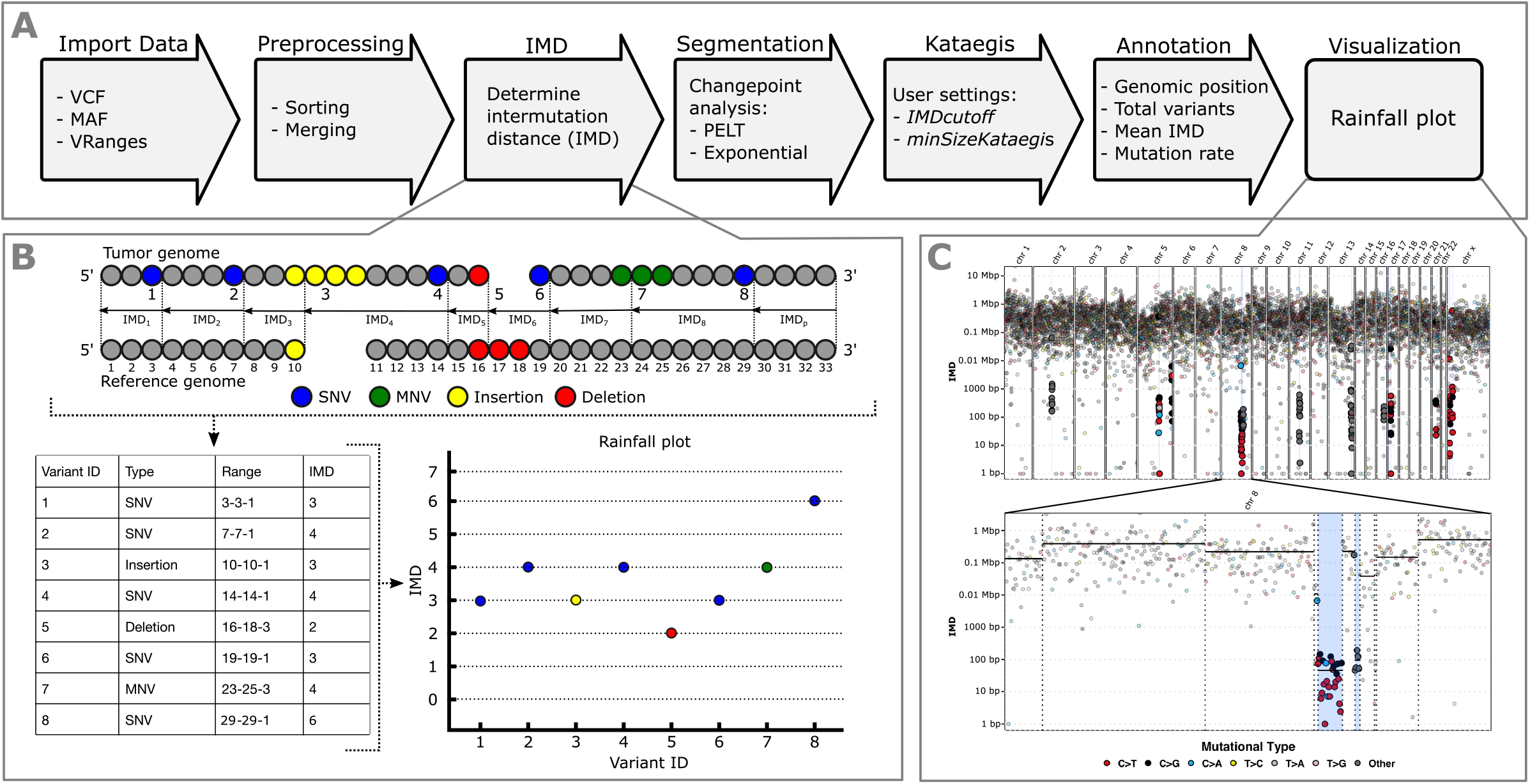
Overview of the Katdetectr workflow, Intermutation distance, and rainfall plots. A) General workflow of Katdetectr from data import to data visualization represented by arrows. B) The intermutation distance (IMD) is determined for all genomic variants in each chromosome, and rainfall plots are used to visualize the IMDs. Single Nucleotide Variant (SNV), Multi Nucleotide Variant (MNV). C) Rainfall plot of WGS breast cancers sample PD7049a as interrogated by Katdetectr with IMDcutoff = 1000 and minSizeKataegis = 6 [1]. Y-axis: IMD, x-axis: variant ID ordered on genomic location, light blue rectangles: kataegis loci with genomic variants within kataegis loci shown in bold. The color depicts the mutational type. The vertical lines represent detected changepoints, while black horizontal solid lines show the mean IMD of each segment.

### Katdetectr search algorithm selection

To optimize Katdetectr for kataegis detection, we generated a synthetic dataset to test four changepoint search algorithms, namely; Pruned Exact Linear Time (PELT) [11], Binary Segmentation (BinSeg) [12], Segment Neighbourhoods (SegNeigh) [13], and At Most One Change (AMOC). The synthetic dataset contains 1024 samples with a varying number of kataegis loci and Tumor Mutational Burden (TMB) (see Methods). All variants in this dataset were binary labeled for kataegis, as a variant either lies within a kataegis locus (TRUE) or not (FALSE). This dataset was considered ground truth and was used for computing performance metrics. We analyzed the synthetic dataset separately for each search algorithm showing that the PELT algorithm outperformed the alternatives (**Supplementary table 1, supplementary figure 1, 2**). Therefore, we set PELT as the default search algorithm in Katdetectr.

### Performance evaluation

We utilized the synthetic dataset to evaluate the performances of Katdetectr and five publicly available kataegis detection packages: MafTools, ClusteredMutations, Kataegis, SeqKat, and, SigProfilerClusters (**Table 1, supplementary table 1**). Katdetectr revealed the highest overall accuracy (0.99), normalized Matthews Correlation Coefficient (nMCC: 0.98), and F1 score (0.97), whereas ClusteredMutations showed the highest True Positive Rate (TPR: 0.99) and Kataegis showed the highest True Negative Rate (TNR: 0.99). Most packages showed a high nMCC for samples with a TMB ranging from 0.1 - 50. However, the performance of all packages dropped for samples with a TMB ≥ 100 (**Figure 2A**). More specifically, for Katdetectr and Kataegis, this is due to an increase in false negatives. For SeqKat, MafTools, ClusteredMutations, and SigProfilerClusters, this performance drop is due to an increase in false positives in samples with a TMB of 100 and 500 (**Supplementary figure 1**).

**Table 1.** Summary information of all evaluated kataegis detection packages and their respective performance metrics regarding kataegis classification on 1024 synthetic samples and 507 a priori labeled Whole Genome Sequenced (WGS) samples. Accuracy, normalized Matthews Correlation Coefficient (nMCC), F1 score, True Positive Rate (TPR) and True Negative Rate (TNR), Pruned Exact Linear Time (PELT), Piecewise Constant Fit (PCF), Intermutation Distance (IMD).

| Package | Reference | Available on | Language | Method | Synthetic dataset |  |  |  |  | WGS dataset |  |  |  |  |
| --- | --- | --- | --- | --- | --- | --- | --- | --- | --- | --- | --- | --- | --- | --- |
|  |  |  |  |  | Accuracy | nMCC | F1 | TPR | TNR | Accuracy | nMCC | F1 | TPR | TNR |
| Katdetectr | Hazelaar, van Riet et al., 2023 | Bioconductor | R | Changepoint analysis (PELT) | 0.99 | 0.98 | 0.97 | 0.94 | 0.99 | 0.99 | 0.92 | 0.83 | 0.91 | 0.99 |
| SeqKat | Taylor et al., 2013 | CRAN | R | Sliding window / exact binomial test | 0.84 | 0.54 | 0.02 | 0.93 | 0.84 | 0.99 | 0.85 | 0.69 | 0.59 | 0.99 |
| MafTools | Mayakonda et al., 2018 | Bioconductor | R | Sliding window / PCF | 0.74 | 0.53 | 0.01 | 0.96 | 0.74 | 0.99 | 0.85 | 0.66 | 0.93 | 0.99 |
| SigProfilerClusters | Bergstrom, Kundu, et al., 2022 | Github | Python | Model sample specific IMD cutoff | 0.65 | 0.52 | 0.01 | 0.88 | 0.65 | 0.99 | 0.84 | 0.68 | 0.66 | 0.99 |
| ClusteredMutations | Lora, 2016 | CRAN | R | Anti-Robinson matrix | 0.70 | 0.53 | 0.01 | 0.99 | 0.74 | 0.99 | 0.83 | 0.61 | 0.99 | 0.99 |
| Kataegis | Lin et al., 2021 | Github | R | PCF | 0.99 | 0.80 | 0.52 | 0.36 | 0.99 | 0.99 | 0.56 | 0.03 | 0.02 | 0.99 |

**Figure 2.**
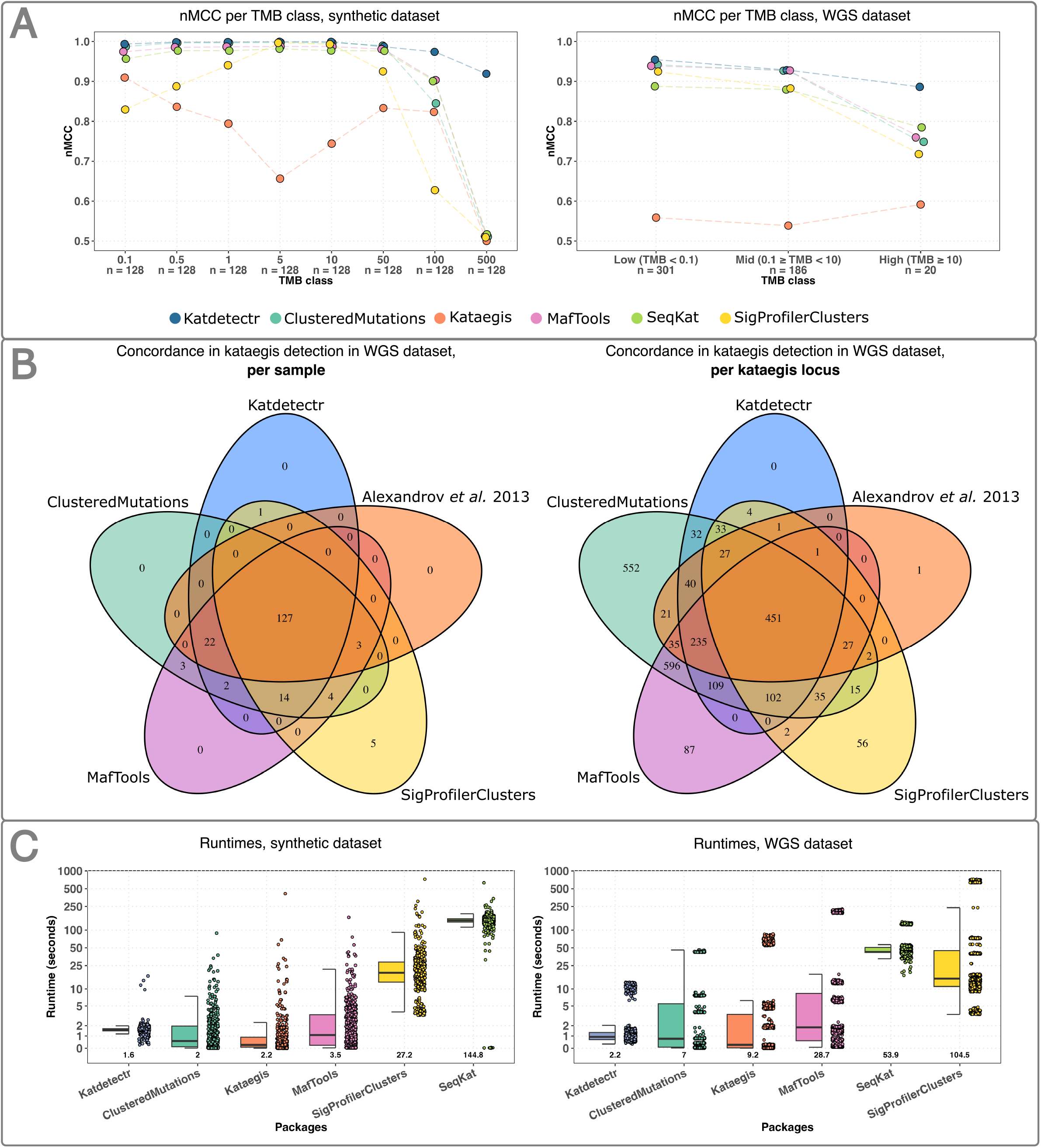
Performance evaluation of kataegis detection tools. A) The normalized Matthews Correlation Coefficient (nMCC) per package and Tumor Mutational Burden (TMB) class is depicted by individual data points connected with a dashed line (colored per package). B) Venn diagrams showing the concordance between Katdetectr, SigProfilerClusters, MafTools, ClusteredMutations, and Alexandrov et al. regarding kataegis classification per sample (i.e., does a sample contain one or more kataegis loci) and per kataegis loci (i.e., does a detected kataegis locus overlap with a kataegis locus detected by another package). C) Boxplots with individual data points represent the per sample runtimes of kataegis detection packages on the synthetic and Whole Genome Sequence datasets. Boxplots were sorted in ascending order based on mean runtime (depicted in the text below the boxplot). Y-axis is log10-scaled. Boxplots depict the Inter Quartile Range, with the median as a black horizontal line.

Next to the synthetic dataset, we evaluated the performance of the kataegis detection packages on a dataset containing 507 *a priori* labeled Whole Genome Sequenced (WGS) samples from Alexandrov *et al*. (see Methods) [1]. Katdetectr revealed the highest overall accuracy (0.99), nMCC (0.92), and F1 score (0.83), whereas ClusteredMutations showed the highest TPR (0.99) and SigProfilerClusters showed the highest TNR (0.99) (**Table 1, Supplementary figure 1)**. Katdetectr, ClusteredMutations, and MafTools showed a high nMCC (>0.92) on the samples with a low or middle TMB. However, the performance of all packages drops for samples with a TMB >10 (**figure 2A**). This is due to an increase in false negatives by Kataegis and SeqKat and false positives by Katdetectr, MafTools, ClusteredMutations, and SigProfilerClusters.

### Summary and performance of kataegis detection packages

We visualized the concordance regarding per sample kataegis classification and kataegis locus between Katdetectr, SigProfilerClusters, ClusteredMutations, MafTools, and the original authors of the WGS dataset: Alexandrov *et al*., *2013* (**Figure 2B**). In total, 451 kataegis loci were detected in 127 WGS samples by all the packages and the original publication. Interestingly, Katdetectr, SigProfilerClusters, ClusteredMutations, and MafTools concordantly detected 102 previously unannotated kataegis loci within the original publication.

The runtimes of all packages were recorded to give insight into the computational feasibility of these packages. Katdetectr showed the lowest mean runtime on both the synthetic and the WGS datasets (**figure 2C**).

### Katdetectr examples with different TMBs

We highlight four samples from the datasets that illustrate how Katdetectr accurately detects kataegis loci regardless of the TMB of the respective sample (**Figure 3**). The synthetic sample 124625_1_50_100 (TMB: 500) harbors one kataegis locus, containing 57 variants, which is detected by Katdetectr (**Figure 3A**). This kataegis locus is also detected by SeqKat, MafTools, ClusteredMutations, and SigProfilerClusters, in addition to numerous false positives. The package Kataegis did not detect any kataegis loci in this synthetic sample.

**Figure 3.**
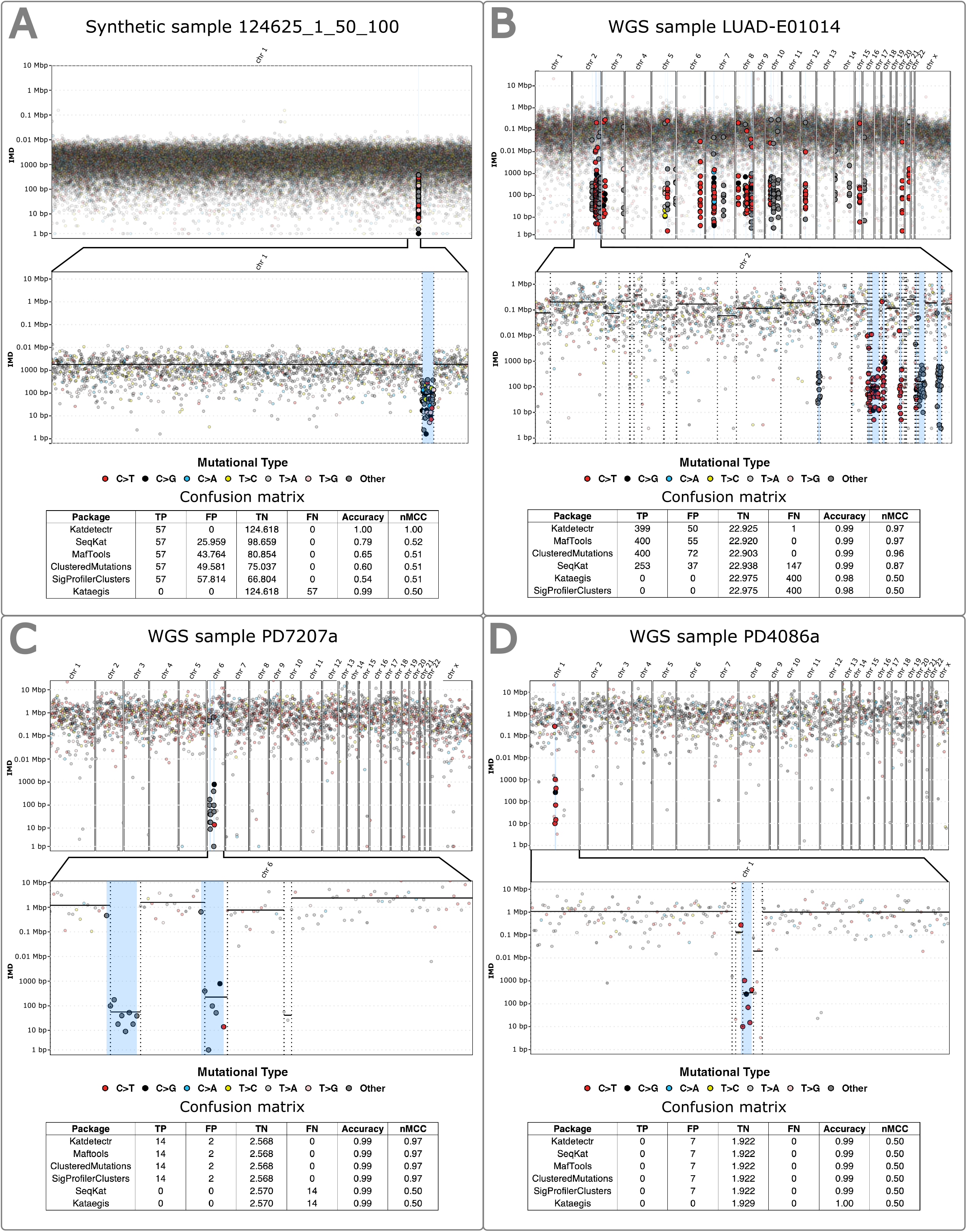
Rainfall plots constructed by Katdetectr and confusion matrices, accuracy, and nMCC for four samples. A) Synthetic sample 124625_1_50_100 with Tumor Mutational Burden (TMB): 500, B) Lung adenocarcinoma Whole Genome Sequenced (WGS) sample LUAD-E01014 with TMB: 7.6. C) Breast cancer WGS sample PD7207a with TMB: 2.5. D) Breast cancer WGS sample PD4086a with TMB: 0.62. The WGS samples were collected and labeled for kataegis by Alexandrov *et al*.; their results were used as ground truth to construct the confusion matrices and performance metrics [1]. Rainfall plot: Y-axis: IMD, x-axis: variant ID ordered on genomic location, light blue rectangles: kataegis loci with genomic variants within kataegis loci shown in bold. The color depicts the mutational type. The vertical lines represent detected changepoints, while black horizontal solid lines show the mean IMD of each segment. Confusion matrix: True Positive (TP), False Positive (FP), True Negative (TN), False Negative (FN), Accuracy, and normalized Matthews Correlation Coefficient (nMCC).

In lung adenocarcinoma sample LUAD-E01014 (TMB: 7.6), Katdetectr detected 37 kataegis loci containing 449 variants (**Figure 3B**). MafTools, ClusteredMutations, and SeqKat detected similar kataegis loci in this sample, whereas Kataegis and SigProfilerClusters did not detect any kataegis loci in this sample. In breast cancer sample PD7207a (TMB: 0.8), two kataegis loci were detected by Katdetectr MafTools, ClusteredMutations, and SigProfilerClusters (**Figure 3C**). Kataegis and SeqKat did not detect any kataegis loci in this sample. Lastly, in the breast cancer sample PD4086a (TMB: 0.6), one kataegis locus was detected by all packages except for Kataegis (**Figure 3D**).

## Discussion

Here, we described Katdetectr, an R/Bioconductor package for the detection, characterization, and visualization of kataegis in genomic variant data by utilizing unsupervised changepoint analysis.

First, we tested four search algorithms for changepoint analysis, which revealed that the PELT [11] algorithm outperformed the BinSeg [12], SegNeigh [13], and AMOC algorithms both in terms of prediction accuracy and computational feasibility. The BinSeg algorithm performed reasonably well, however, it underfitted the data, which resulted in many false negatives. The SegNeigh algorithm performed well on samples with a TMB < 5; however, this algorithm is computationally expensive, as it scales exponentially with the size of the data, and cannot reasonably be used for the analysis of samples with a TMB > 10. Unsurprisingly, the AMOC (at most one change) algorithm cannot detect kataegis as a kataegis locus is generally defined by two changepoints.

Besides testing search algorithms, we benchmarked Katdetectr using PELT and five publicly available kataegis detection packages which were recently published and used for supporting kataegis research [2, 5, 14, 15]. Since no consensus benchmark was available, we aimed to get insight into the performance of these tools. The complexity of kataegis detection is to separate genomic regions of higher-than-expected mutational density from the background of somatic mutations. Therefore, we argued that generating a synthetic dataset containing samples of varying TMB (0.1-500), would provide a good measure for algorithmic solvability of the kataegis detection problem. Benchmarking on this synthetic dataset revealed that the accuracy of kataegis detection for all evaluated packages drops when the TMB increases. Performance evaluation per TMB-binned class revealed that Katdetectr is on par with alternative packages for samples with low or middle TMB. However, in contrast to alternative packages, Katdetectr remained robust when analyzing samples with a high TMB. Additionally, the computation times of Katdetectr are feasible for samples with a TMB ranging from 0.1 to 500 as PELT scales linearly with the size of the data [11]. This shows that kataegis detection using Katdetectr is feasible on reasonably modern computer hardware.

The presented performance evaluation depends on the truth labels provided by the datasets. Both the synthetic and the WGS dataset have their limitations. We constructed the synthetic dataset by modeling mutations on a genome as a Bernoulli process, which is a common approach for modeling events that occur in a sequence. However, we did not incorporate prior biological knowledge in the synthetic dataset generation. Both SeqKat and SigProfilerClusters incorporate biological assumptions regarding kataegis, e.g., mutation context, which possibly negatively influenced their performance regarding the synthetic dataset. Additionally, the distance between events generated by a Bernoulli process is a geometric random variable. For a large n, which is the case for a human genome, a geometric random variable approximates an exponential random variable. Since we constrain Katdetectr to only fit exponential distributions it is unsurprising that Katdetectr performs well on the synthetic dataset. Nevertheless, MafTools, ClusteredMutations, SeqKat, and SigProfilerClusters are less robust when analyzing the synthetic samples with a TMB of 100 and 500 as they classify many false positives kataegis loci.

In addition to the synthetic dataset, we used the *a priori* labeled pan-cancer WGS dataset from the groundbreaking work of Alexandrov *et al*. to evaluate the kataegis detection tools [1]. However, the field of kataegis has grown and evolved since the publication of this dataset. Therefore, we want to emphasize that this dataset should not be considered an unequivocal truth, and the performance metrics should not be taken at face value. The annotation of this dataset likely contains several false positives and false negatives; as highlighted by the concordant discovery of 102 additional kataegis foci by several packages. Nevertheless, as no alternative ground-truth has been established to determine the accuracy of kataegis detection, we believe that the current benchmarking results give insight into the behavior of the evaluated packages regarding kataegis classification in samples with varying TMB.

Our benchmarking showed that, for the WGS dataset, Katdetectr, MafTools, ClusteredMutations, and, SigProfilerClusters have a high concordance in classifying a whole sample as kataegis positive or negative. However, when concerning distinct kataegis loci, we observed more differences. ClusteredMutations reported the overall largest number of loci (n = 2,360), indicating it has the highest sensitivity. Conversely, kataegis (n = 8) and SeqKat (n = 528) reported the overall smallest number of loci which we deem too small based on visual inspection. The third smallest number of kataegis loci is reported by SigProfilerClusters (n = 764), indicating it has the highest specificity. Katdetectr appears to balance sensitivity and specificity as it only detects kataegis loci detected by one or more alternative packages (n = 1,050).

Kataegis is the most commonly used term for local hypermutations and has historically been defined as a cluster of at least six variants, of which the mean IMD is less or equal to 1000 base pairs [1, 16]. However, this definition has been altered recently, making the formal definition of kataegis ambiguous [2, 4, 5, 14]. For instance, another type of clustered mutations is called Omikli, which refers to clusters smaller than kataegis, generally containing three or four variants [17]. Although different types of clustered variants can be detected using Katdetectr by supplying the correct parameters, we only evaluated Katdetectr for the detection of kataegis.

We made Katdetectr publicly available on the Bioconductor platform, which requires peer-reviewed open-source software and high standards regarding development, documentation, and unit testing. Furthermore, Bioconductor ensures reliability and operability on common operating systems (Windows, macOS, and Linux). We designed Katdetectr to fit well in the Bioconductor ecosystem by incorporating common Bioconductor object classes. This allows Katdetectr to be used reciprocally with the plethora of statistical software packages available in Bioconductor for preprocessing and subsequent analysis. Lastly, we implemented Katdetectr flexibly, allowing Katdetectr to be used in an ad hoc manner for quick assessment of clustered variants and extensive research of the mutation rates across a tumor genome.

## Conclusion

Katdetectr is a free, open-source R package available on Bioconductor that contains a suite for the detection, characterization, and visualization of kataegis. Katdetectr employs the PELT search algorithm for unsupervised changepoint analysis, resulting in robust and fast kataegis detection. Additionally, Katdetectr has been implemented in a flexible manner which allows Katdetectr to expand in the field of kataegis. Katdetectr is available on Bioconductor: https://doi.org/doi:10.18129/B9.bioc.katdetectr and on GitHub: https://github.com/ErasmusMC-CCBC/katdetectr.

## Acknowledgments

We thank Martijn Lolkema, John Martens, Marcel Smid, Guido Jenster, and Stavros Makrodimitris for their discussions, input, and support. Additionally, we would like to thank Coen Berns and Yi Ping for their initial efforts in detecting kataegis.

## Methods

### Implementation Katdetectr

Katdetectr (v1.1.3, git commit 8cf59018956b6d724f3b66d03b343011ad24e554) was developed in the R statistical programming language (v4.2.0) [18]. Katdetectr imports genomic variants through generic, standardized file formats for variant calling: MAF, VCF, or Bioconductor-standard VRanges objects. Within Katdetectr, the imported variants are pre-processed such that, per chromosome, all variants are sorted in ascending order based on their genomic position. Overlapping variants are merged into a single record. Following, per *chromosome*_*j*_, the intermutation distance (*IMD*_*i,j*_) of each *v*ariant_*i,j*_ and its closest upstream *variant*_*i* − 1,*j*_ is calculated according to;

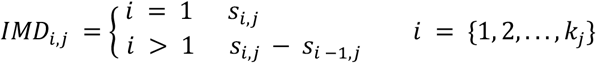

With *i* as the variant number, *j* as the chromosome number, *s* as the genomic location of the first base-pair of a *variant*_*i,j*_ and *k*_*j*_ as the total number of variants in *chromosome*_*j*_ (**Figure 1B**). Additionally, for each *chromosome*_*j*_ one pseudo IMD, *IMD*_*p,j*_, is added such that;

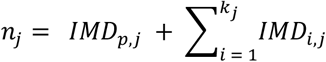

With *n*_*j*_ as the total number of base-pairs in *chromosome*_*j*_.

Katdetectr aims to identify genomic regions characterized by specific mutation rates. An unsupervised technique called changepoint analysis is performed per chromosome on the IMDs to assess the variability in mutation rate across each chromosome. Changepoint analysis refers to the process of detecting points in a sequence of observations where the statistical properties of the sequence significantly change. Subsequently, the detected changepoints are used to segment the input sequence into segments. For a detailed description of the changepoint analysis, see the work of Killick, Fearnhead, and Eckley [11]

We implemented the commonly used R changepoint package (v2.2.3) in Katdetectr for the unsupervised segmentation of IMDs, as detailed by [11, 20, 21]. We set the following parameters settings; method: Pruned Exact Linear Time (PELT), minimal segment length: 2, test statistic: Exponential, and penalty: Bayesian Information Criterion (BIC), as default settings in Katdetectr.

After changepoint analysis, each segment is annotated with its respective genomic start and end positions, its mean IMD, and the total number of included variants. Since we use an exponential distribution as the test statistic in changepoint analysis, each segment has a corresponding rate parameter of the fitted exponential distribution. Whereas each segment is annotated with its corresponding mutation rate, the mutation rate of an entire sample can be expressed as the weighted arithmetic mean of the mutation rate of the segments;

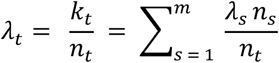

With *λ*_*t*_ as the mutation rate of the entire sample, *k*_*t*_ as the total number of variants present in the sample, *n*_*t*_ as the total number of base pairs in the genome, *m* as the total number of segments in the sample, and *λ*_*s*_ and *n*_*s*_ as the mutation rate and the number of base-pairs in *segment*_*s*_

To call a segment a putative kataegis locus, it has to adhere to two user-defined parameters: the maximum mean IMD of the segment (*IMDcutoff*) and the minimum number of included variants (*minSizeKataegis*). These parameters can be provided as static integer values or as a custom R function determining the IMD cutoff for each segment. For example, the following function for annotation of kataegis events, as was used by the ICGC/TCGA Pan-Cancer Analysis of Whole Genomes Consortium, can be easily implemented in Katdetectr [4]:

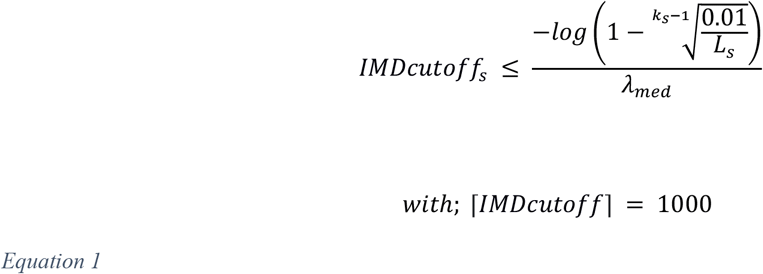

With *IMDcutoff*_*s*_ as the IMD cut-off value, *k*_*s*_ as the number of mutations and *L*_*s*_ as the length of *segment*_*s*_ in base-pairs. For this function the rate of the whole sample is modeled assuming an exponential distribution with;

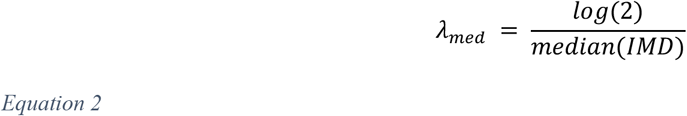

Henceforth, all segments satisfying these user-specified parameters are considered putative kataegis loci and stored appropriately. Two or more adjacent kataegis loci are merged and stored as a single record.

The output of Katdetectr consists of an S4 object of class “KatDetect” which stores all relevant information regarding kataegis detection and characterization. A KatDetect object contains four slots: 1) the putative kataegis loci (Granges), 2) the detected segments (Granges), 3) the inputted genomic variants with annotation (Vranges), and 4) the parameters settings (List). These data objects can be accessed using accessor functions.

In addition, we implemented three methods for the KatDetect class, *summary, show*, and *rainfallPlot*. In concordance with R standards, the *summary* function prints a synopsis of the performed analysis, including the number of detected kataegis loci; and the number of variants inside a kataegis loci. The *show* function displays information regarding the S4 class and the synopsis.

The method *rainfallPlot* is a function for generating rainfall plots. These rainfall plots display the genomic ordered IMDs (from all genomic variants) within a sample and highlight putative kataegis loci and associated genomic variants. This function has additional arguments: *showSequence*, which allow the user to display specific chromosomes, and *showSegmentation*, for displaying the changepoints and the mean IMD of all segments.

For additional examples and more hands-on technical instructions, we refer to the accompanying vignette (**Supplemental vignette**) or the online Bioconductor repository (https://bioconductor.org/packages/release/bioc/vignettes/katdetectr/inst/doc/General_overview.html).

### Performance evaluation

As multiple packages for kataegis detection are publicly available, we compared Katdetectr against MafTools (v2.13.0), ClusteredMutations (v1.0.1), kataegis (v0.99.2), SeqKat (v0.0.8) and, SigProfilerClusters (v1.0.11) [6-10]. For benchmarking, we used an in-house generated synthetic dataset and an *a priori* labeled pan-cancer dataset of whole genome sequenced malignancies.

We used the following definition of kataegis as postulated by Alexandrov and colleagues: a kataegis locus is 1) a continuous segment harboring ≥6 variants and 2) the captured IMDs within the segment have a mean IMD of ≤1000 bp [1]. To quantify and compare performances, the task of kataegis detection was reduced to a binary classification problem. The task of the kataegis detection packages was to correctly label each variant for kataegis, i.e., whether or not a genomic variant lies within a kataegis locus.

### Performance metrics

Only a small fraction of all observed variants is located within kataegis loci, this results in a large class imbalance which renders the interpretation of performance metrics, such as accuracy, F1, TPR, and TNR, counterintuitive and possibly unrepresentative (**Equation 3**). Therefore, the normalized Matthews Correlation Coefficient (nMCC) was used as the primary metric for performance evaluation. The nMCC considers performance proportionally to both the size of positive and negative elements in a dataset [21].

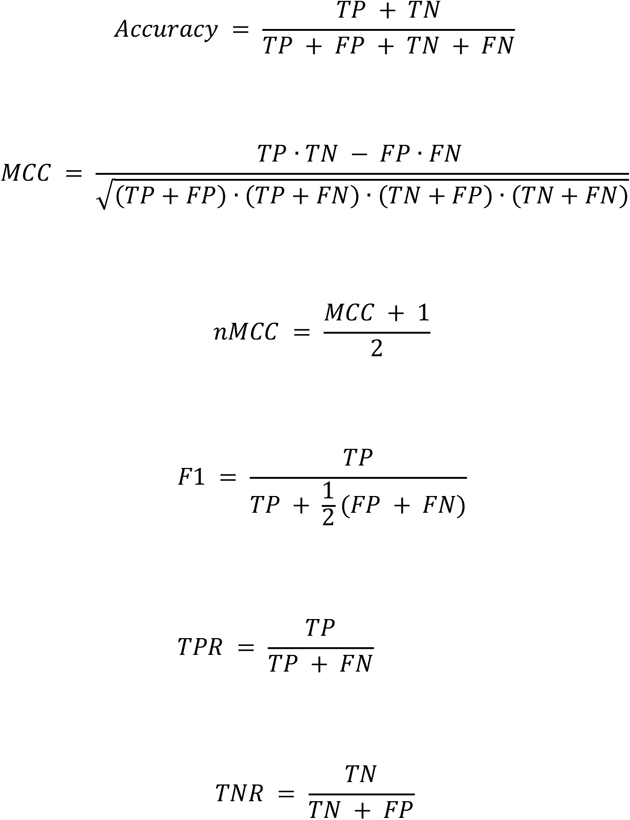

*Equation 3. Performance metrics. Accuracy, Matthews Correlation Coefficient (MCC), normalized Matthews Correlation Coefficient (nMCC), F1 score, True Positive Rate (TPR), and True Negative Rate (TNR)*.

*True Positive (TP): Predicted: variant in kataegis locus. Truth set: variant in kataegis locus*.

*False Positive (FP): Predicted: variant in kataegis locus. Truth set: variant not in kataegis locus*.

*True Negative (TN): Predicted: variant not in kataegis locus. Truth set: variant not in kataegis locus*.

*False Negative (FN): Predicted: variant not in kataegis locus. Truth set: variant in kataegis locus*.

We utilized Venn diagrams to display the concordance of the kataegis detection packages. We showed in which samples the packages detected one or more kataegis loci and which kataegis loci were detected by the packages. Two packages are said to detect the same kataegis locus if the genomic locations of their respective kataegis locus overlap by at least one base pair.

To give insight into the packages computation time, the packages runtime performance was recorded using the `proc.time()` function from the base R package. All packages and comparisons were run on the same server utilizing an AMD EPYC 7742 64-Core Processor. The packages Katdetectr and SigProfilerClusters contained options for parallel processing and used at most four cores per sample during the analyses. All other packages used a single processing core per sample.

All scripts necessary for running and visualizing the performance evaluation of all evaluated packages are available on GitHub at https://github.com/ErasmusMC-CCBC/evaluation_katdetectr. All data used for the performance evaluation is available at https://dx.doi.org/10.5281/zenodo.6810477.

### Synthetic data generation

The synthetic dataset was generated using the `generateSyntheticData()` function within the Katdetectr package. Mutations were randomly sampled on a reference genome such that each base has an equal probability, *p*, of being mutated (except for N bases for which *p* = 0). This reduces the occurrence of mutations on the reference genome to a sequence of *X*_*1*_, *X*_*2*_, …, *X*_*n*_, independent Bernoulli trials, *X*_*i*_, i.e., a Bernoulli process, where;

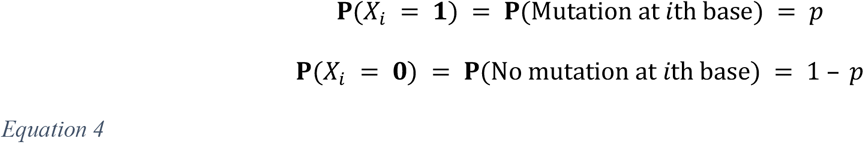

with probability mass function (PMF), expectation and variance:

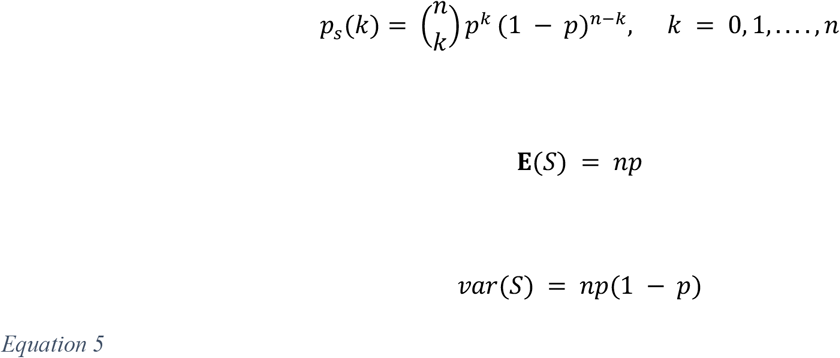

with *p* as the probability of success (i.e., mutation), *n* as the number of independent trials (i.e., length of the genome in base pairs), and *k* as the number of successes (i.e., number of occurred mutations). The IMD now reduces to geometric random variable *T*; with PMF, expectation, and variance:

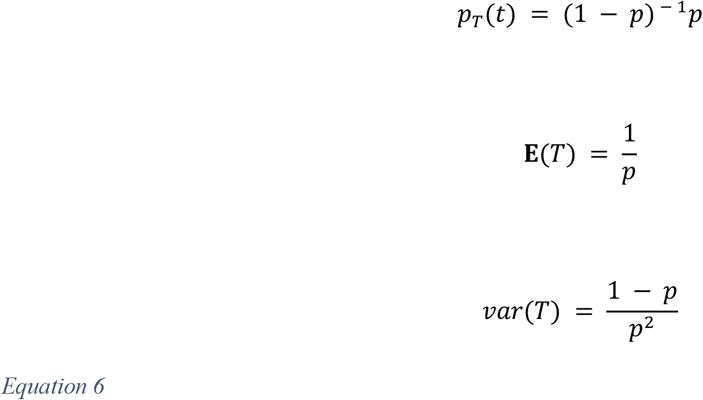

The genomic start location of a kataegis locus was sampled as an independent Bernoulli trial. The genomic end location of a kataegis locus was calculated using:

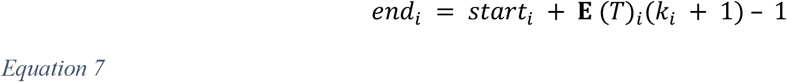

### Synthetic dataset description

The synthetic data consists of 1024 samples with a total of 21.299.360 SNVs (**Table 2**). All mutations were generated on chromosome 1 on the human reference genome hg19. These samples were generated such that 8 different TMB classes (0.1, 0.5, 1, 5, 10, 50, 100, 500) were considered.

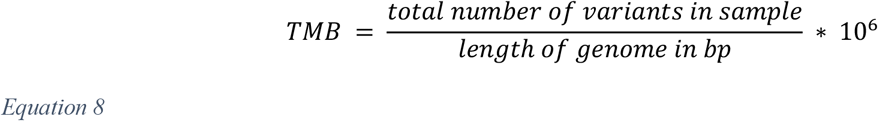

For each TMB class, a sample was generated for all combinations of the following parameters: the number of kataegis loci (1, 2, 3, 5); the number of variants within each kataegis loci (6, 10, 25, 50); and the expected IMD of the variants in kataegis loci (100, 250, 500, 750). This resulted in 64 kataegis samples per TMB class. To balance the dataset, 64 samples without kataegis loci were generated for each TMB class. The synthetic dataset contained 1232 kataegis loci and 33.245 variants within kataegis loci.

### Descriptive statistics of synthetic dataset

**Table 2.** Showing per Tumor Mutational Burden (TMB) class: TMB, number of generated background mutations per sample, the total number of samples, total number of samples with kataegis, total number of kataegis loci, and total number of variants within a kataegis loci of 1024 synthetic samples.

| TMB class (no. of background mutations) | No. Samples (with kataegis) | No. Kataegis loci | No. Variants in kataegis loci |
| --- | --- | --- | --- |
| 0.1 (25) | 128 (64) | 176 | 4.005 |
| 0.5 (125) | 128 (64) | 176 | 4.006 |
| 1 (249) | 128 (64) | 176 | 4.006 |
| 5 (1.246) | 128 (64) | 176 | 4.014 |
| 10 (2.493) | 128 (64) | 176 | 4.029 |
| 50 (12.463) | 128 (64) | 176 | 4.077 |
| 100 (24.925) | 128 (64) | 176 | 4.183 |
| 500 (124.625) | 128 (64) | 176 | 4.925 |

### Whole Genome Sequence (WGS) dataset description

The WGS dataset (as used in this study; **table 3**) is publicly available in .txt format[1]. This dataset contained 7.042 primary cancer samples from 30 different tissues; of which 507 originate from whole genome sequencing (WGS) and 6.535 from whole exome sequencing (WES). Only the WGS samples (*n* = 507) were originally labeled using a Piece-Wise Constant Fit (PCF) model and manually curated for kataegis presence (or absence) by the original study. Only the respective WGS samples were re-interrogated within our performance evaluation. Additionally, we binned this dataset into three TMB classes (low: TMB < 0.1, middle: 0.1 ≥ TMB < 10, high: TMB ≥ 10) and filtered it such that it only contained single nucleotide variants (SNVs).

### Descriptive statistics of WGS dataset

**Table 2.** Showing per Tumor Mutational Burden (TMB) class: TMB range, the total number of samples, total number of samples with kataegis, total number of kataegis loci, and total number of variants within a kataegis loci of 507 Whole Genome Sequenced (WGS) samples labeled by Alexandrov *et al*. [1].

| TMB class | # Samples (with kataegis) | # Kataegis loci | # Variants in kataegis loci |
| --- | --- | --- | --- |
| Low: TMB < 0.1 | 301(45) | 93 | 946 |
| Middle: $0.1 \geq \text{TMB} < 10$ | 186 (89) | 444 | 5.058 |
| High: TMB $\geq 10$ | 20(18) | 336 | 3.107 |

### Pre-processing and parameter settings of alternative kataegis detection packages

Both the synthetic and the Alexandrov et al. datasets were converted to MAF format for use in MafTools [6] ClusteredMutations [7], and kataegis [8] and to BED format for use in SeqKat [9]. All other parameter settings for MafTools, kataegis, ClusteredMutations, and SeqKat were set to the default values as specified in their respective manuals and vignettes.

For SigProfilerClusters [10] both the synthetic and the Alexandrov et al. datasets were converted to a .txt file with column names as specified in the manual of SigProfilerClusters. We set the following parameters for SigProfilerSimulator(): genome="GRCh37", contexts = [‘288’], simulations=100, overlap=True. For subsequent cluster detection, we set the following parameters for SigProfilerClusters.analysis(): genome="GRCh37", contexts="96", simContext=["288"], analysis="all", sortSims=True, subClassify=True, correction=True, calculateIMD=True, max_cpu=4, includedVAFs=False.

From the output of SigProfilerClusters we selected the class 2 (kataegis) clusters for further analysis. The definition of kataegis used by SigProfilerClusters differs from the one used in our performance evaluation. SigProfilerClusters defines kataegis as a cluster of ≥4 genomic variants of which the mean IMD is statistically different from the sample specific IMD cut-off. To include SigProfilerClusters in our performance evaluation we only selected clusters detected by SigProfilerClusters that fit the definition of kataegis we used for the performance evaluation, i.e., a kataegis locus contains ≥6 genomic variants with a mean IMD ≤1000 bp.

## Funding

This research received funding from the Daniel den Hoed Fonds - Cancer Computational Biology Center (DDHF-CCBC) grant.

## Conflict of Interest

None declared.

## Data availability

All data used in the performance evaluation can be found on Zenodo at: https://dx.doi.org/10.5281/zenodo.6810477.

## List of abbreviations

AMOC: At Most One Change
bp: base-pair
BinSeg: Binary Segmentation
IMD: Intermutation Distance
MAF: Mutation Annotation Format
MNV: Multi Nucleotide Variant
nMCC: normalized Matthews Correlation Coefficient
PCF: Piecewise Constant Fit
PELT: Pruned Exact Linear Time
SNV: Single Nucleotide Variant
SegNeigh: Segment Neighbourhoods
TMB: Tumor Mutational Burden
TNR: True Negative Rate
TPR: True Positive Rate
VCF: Variant Calling Format
WES: Whole Exome Sequencing
WGS: Whole Genome Sequencing

## Availability and requirements

- Project name: **Katdetectr**
- Project home page:
  - https://bioconductor.org/packages/release/bioc/html/katdetectr.html
  - https://github.com/ErasmusMC-CCBC/katdetectr
- Operating system(s): Platform independent
- Programming language: R (>= 4.2)
- Other requirements: BiocParallel (>= 1.26.2), changepoint (>=2.2.3), checkmate (>= 2.0.0), dplyr (>= 1.0.8), GenomicRanges (>= 1.44.0), GenomeInfoDb (>= 1.28.4), IRanges (>=2.26.0), maftools (>= 2.10.5), methods (>= 4.1.3), rlang (>= 1.0.2), S4Vectors (>= 0.30.2), tibble (>= 3.1.6), VariantAnnotation (>= 1.38.0), Biobase (>= 2.54.0), Rdpack (>= 2.3.1), ggplot2 (>= 3.3.5), tidyr (>= 1.2.0), BSgenome (>= 1.62.0), ggtext (>= 0.1.1), BSgenome.Hsapiens.UCSC.hg19 (>= 1.4.3), BSgenome.Hsapiens.UCSC.hg38 (>= 1.4.4), plyranges (>= 1.17.0)
- License: GPL-3
- Project name: **Evaluation of Katdetectr and alternative kataegis detection packages**
- Project home page: https://github.com/ErasmusMC-CCBC/evaluation_katdetectr
- Operating system(s): Platform independent
- Programming language: R (>= 4.2)
- Other requirements: katdetectr (1.1.2), MafTools (2.13.0), ClusteredMutations (1.0.1), kataegis (0.99.2), SeqKat (0.0.8), SigProfilerClusters (1.0.11), dplyr (1.0.10), tidyr (1.2.1), ggplot2 (3.4.0), variantAnnotation (1.44.0), mltools (0.3.5)
- License: GPL-3

## Author contributions

Daan M. Hazelaar: Conceptualization, Data curation, Formal Analysis, Investigation, Methodology, Software, Validation, Visualization, Writing – Original draft

Job van Riet: Conceptualization, Methodology, Investigation, Software, Visualization, Writing – review & editing

Youri Hoogstrate: Conceptualization, Methodology, Software, Writing - review & editing

Harmen J. G. van de Werken: Conceptualization, Funding acquisition, Investigation, Methodology, Project administation, Resources, Supervision, Writing - review & editing

## Notes

https://www.bioconductor.org/packages/devel/bioc/html/katdetectr.html

https://github.com/ErasmusMC-CCBC/evaluation_katdetectr

https://zenodo.org/record/6623289#.YqBxHi8Rr0o

### Competing Interest Statement

The authors have declared no competing interest.

### Summary of Updates

Extended the performance evaluation.

